# Discordant effects of *ex-vivo* JAK inhibition on inflammatory responses in colonic compared to ileal mucosa

**DOI:** 10.1101/2024.01.18.576013

**Authors:** Kawsar Kaboub, Hanan Abu-Taha, Jessica Arrouasse, Efrat Shaham-Barda, Nir Wasserberg, Lucille Hayman-Manzur, Adi Friedenberg, Adva Levy-Barda, Idan Goren, Henit Yanai, Zohar Levi, Hagar Banai-Eran, Irit Avni-Biron, Jacob E Ollech, Sarit Cohen-Kedar, Iris Dotan, Keren M Rabinowitz

## Abstract

**Background & aims:** Janus kinase (JAK) inhibitors modulating JAK-STAT (signal transducers and activators of transcription) signaling pathway, are used for the treatment of patients with inflammatory bowel diseases (IBD). We aimed to identify the molecular effects of JAK inhibition in the human intestinal mucosa, considering the IBD location and phenotype.

**Methods:** Colonic and ileal explants from patients with ulcerative colitis (UC), Crohn’s disease (CD), or non-IBD controls (NC) were treated *ex-vivo* with the JAK inhibitor, tofacitinib. Phosphorylated STAT (p-STAT) levels were assessed by Western blot and Immunofluorescence. Inflammatory genes expression was assessed with Nanostring nCounter system. Human intestinal organoids were used to assess JAK inhibitors’ effects on p-STATs and iNOS expression.

**Results:** Explants were collected from 68 patients (NC=28; UC=20; CD=20). JAK inhibition reduced p-STAT1/3/5 expression in all explants. While p-STAT inhibition rates varied among patients (10%-88%), higher inhibition rates were observed in colonic compared to ileal explants. Significant alterations in 120 of 255 inflammatory genes were observed in colonic explants, while only 30 were observed in ileal NC explants. In colonic explants from UC, significant alterations were observed in 5 genes, including *STAT1* and *NOS2*. Various JAK inhibitors reduced IFN-γ-induced increase in p-STAT1 and iNOS expression in organoids.

**Conclusions:** A site-specific anti-inflammatory effect of JAK inhibition by tofacitinib was noticed, whereby the colon was more robustly affected than the ileum. *Ex-vivo* response to tofacitinib is individual. JAK inhibition may attenuate inflammation by decreasing iNOS expression. *Ex-vivo* mucosal platforms may be a valuable resource for studying drug impact and evaluating personalized treatment effects.

## Introduction

Crohn’s Disease (CD) and ulcerative Colitis (UC), the two main forms of inflammatory bowel diseases (IBD), are characterized by chronic remitting inflammation. CD involves inflammation that may affect any part of the gastrointestinal tract, while inflammation in UC is restricted to the colon^1,2^. The orally administered small molecule Janus kinase (JAK) inhibitors, used for the treatment of IBD^3,4^, block the JAK-signal transducer and activator of transcription (STAT) pathways responsible for transmitting pro-inflammatory cytokines in IBD, including interferon (IFN)-γ, IL-6, IL-9, and IL-12/23^5–7^.

The JAK family comprises four intracellular non-receptor tyrosine kinases (TYKs), including JAK1, JAK2, JAK3, and TYK2^8^. Multiple JAK inhibitors with various specificities have been developed for the treatment of IBD, demonstrating different efficacies in UC and CD. Tofacitinib, a pan-JAK inhibitor, is used for treating moderate to severe active UC. However, in CD trials remission rates in patients receiving tofacitinib were comparable to placebo^9,10^. In contrast, the selective JAK1 upadacitinib, exhibits efficacy and is used for treating both UC and CD^11,12^.

JAK inhibitors’ mechanism of action in the inflamed human intestinal mucosa, is still elusive. Moreover, the factors contributing to the differential efficacy of JAK inhibitors in patients with UC and CD are still unclear. Here, we investigated the mechanism of action of JAK inhibition in the intestinal mucosa extending to various intestinal locations and IBD phenotypes. To this end, we used fresh human colonic and ileal mucosa from non-IBD controls (NC) and patients with UC or CD to study the effects of JAK inhibition on signaling, inflammatory genes expression, and intestinal cellular function *ex-vivo*.

## Material and Methods

### Patients and tissues

IBD colonic and ileal mucosal explants were obtained from surgical specimens of patients with IBD undergoing colonic or ileal resection, or from biopsies of patients undergoing routine colonoscopies at the Rabin Medical Center. Non-IBD colonic and ileal mucosal explants were obtained from surgical specimens of patients undergoing bowel resection for colonic tumors (normal mucosa taken from a distance of at least 10 cm from the tumor), or from biopsies of patients undergoing colonoscopy for colon-cancer screening, diverticulitis, or irritable bowel syndrome. The study included 68 patients; 26 surgical specimens were used for p-STATs levels assessment and 42 biopsies were used for inflammatory gene expression levels assessment. Patients’ clinical and demographic characteristics are detailed in Table 1.

**Table 1.**
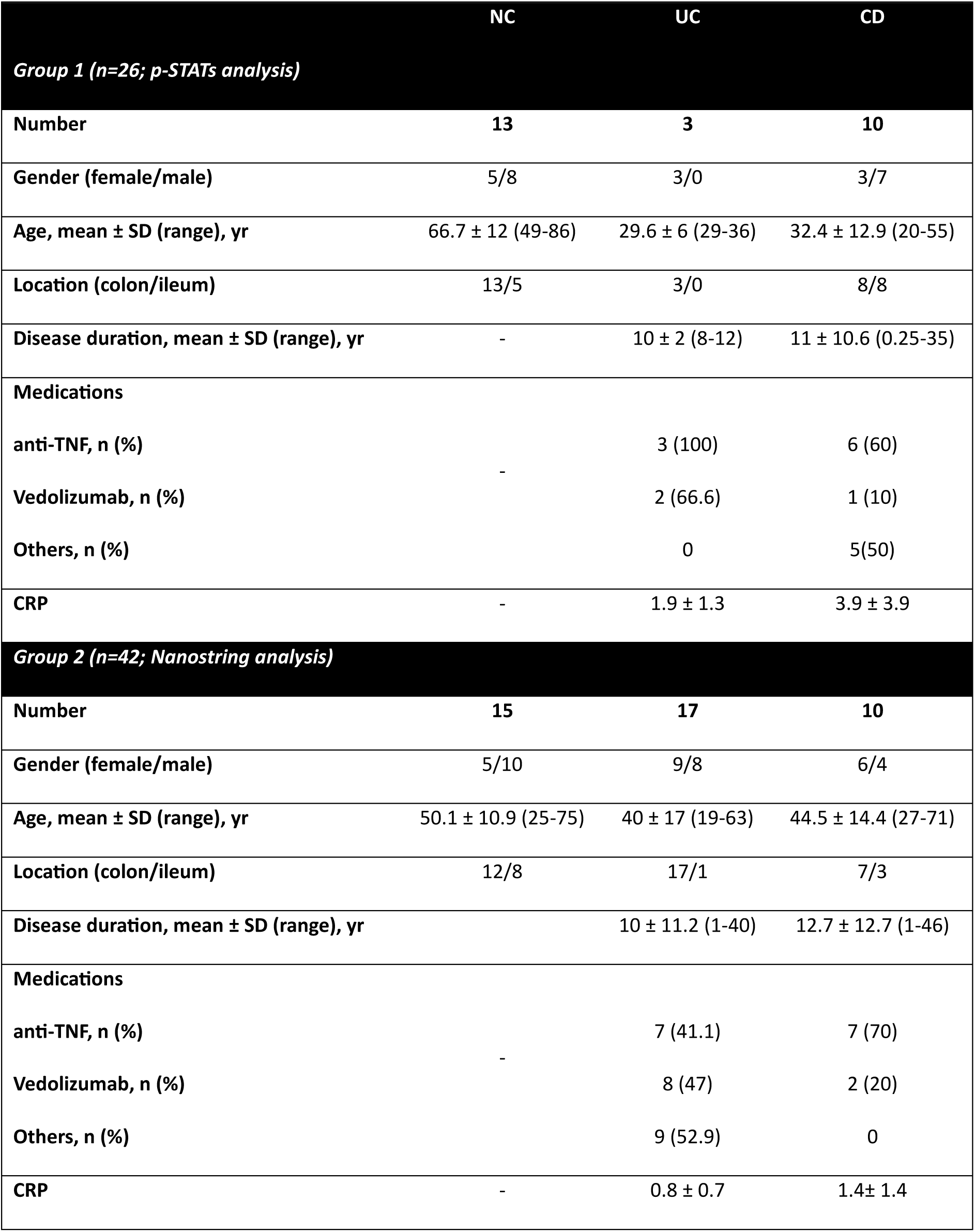
Clinical and demographic characteristics of patients included in the study. NC- non-IBD control; UC- ulcerative colitis; CD- Crohn’s disease; p-STAT- phosphorylated signal transducer and activator of transcription; TNF- Tumor necrosis factor; CRP- C- reactive protein.

|  | NC | UC | CD |
| --- | --- | --- | --- |
| **Group 1 (n=26; p-STATs analysis)** |  |  |  |
| **Number** | **13** | **3** | **10** |
| **Gender (female/male)** | 5/8 | 3/0 | 3/7 |
| **Age, mean ± SD (range), yr** | 66.7 ± 12 (49-86) | 29.6 ± 6 (29-36) | 32.4 ± 12.9 (20-55) |
| **Location (colon/ileum)** | 13/5 | 3/0 | 8/8 |
| **Disease duration, mean ± SD (range), yr** | - | 10 ± 2 (8-12) | 11 ± 10.6 (0.25-35) |
| **Medications** |  |  |  |
| **anti-TNF, n (%)** | - | 3 (100) | 6 (60) |
| **Vedolizumab, n (%)** | - | 2 (66.6) | 1 (10) |
| **Others, n (%)** | - | 0 | 5(50) |
| **CRP** | - | 1.9 ± 1.3 | 3.9 ± 3.9 |
| **Group 2 (n=42; Nanostring analysis)** |  |  |  |
| **Number** | **15** | **17** | **10** |
| **Gender (female/male)** | 5/10 | 9/8 | 6/4 |
| **Age, mean ± SD (range), yr** | 50.1 ± 10.9 (25-75) | 40 ± 17 (19-63) | 44.5 ± 14.4 (27-71) |
| **Location (colon/ileum)** | 12/8 | 17/1 | 7/3 |
| **Disease duration, mean ± SD (range), yr** | - | 10 ± 11.2 (1-40) | 12.7 ± 12.7 (1-46) |
| **Medications** |  |  |  |
| **anti-TNF, n (%)** | - | 7 (41.1) | 7 (70) |
| **Vedolizumab, n (%)** | - | 8 (47) | 2 (20) |
| **Others, n (%)** | - | 9 (52.9) | 0 |
| **CRP** | - | 0.8 ± 0.7 | 1.4 ± 1.4 |

The Institutional Ethical Committee of the Rabin Medical Center approved the study (approval number 0547-17-RMC, 0763-16-RMC and 0298-17). A written informed consent of all participating subjects was obtained.

### Intestinal mucosal explants culture *ex-vivo*

Intestinal mucosal explants were immediately transferred from endoscopy suit to the research laboratory in complete medium, consisting of RPMI 1640 (01-100-1A, Biological industries, Beit Haemek, Israel) supplemented with 10% fetal bovine serum (FBS) (03-007-1A, Biological industries, Beit Haemek, Israel), 100 U/ml penicillin and 100 µg/ml streptomycin (03-031-1B, Biological industries, Beit Haemek, Israel), and 2.5 µg/ml amphotericin B (Fungizone, 03-028-1B, Biological industries, Beit Haemek, Israel) on ice. Mucosa was washed with sterile phosphate buffer saline (PBS), stripped from the resected tissue, and cut into 2– 3 mm^3^ explants. Mucosal explants were cultured in a complete medium supplemented with 100µg/ml gentamicin (03-035-1B, Biological Industries, Beit Haemek, Israel) in a 37°C 5% CO2 incubator. For p-STATs assessment, mucosal explants were treated with 10-1000 nM tofacitinib (PZ0017, Sigma-Aldrich, Missouri, United States) for 1.5h. For inflammatory genes expression assessment mucosal explants were treated with 10 µM tofacitinib or 0.1% DMSO (67-68-5, Sigma-Aldrich, Missouri, United States) control (CTRL, the concentration used as tofacitinib’s vehicle) overnight. 0.1% of vehicle DMSO did not affect p-STATs levels, thus it was used as CTRL (Fig 1A). Subsequently, mucosal explants were snap frozen in either liquid nitrogen or Tissue–Plus OCTcompound (Scigen Scientific Gardena, California, United States) and stored at −80°C until protein or RNA extraction, or sectioning for immunofluorescence, respectively.

**Figure 1.**
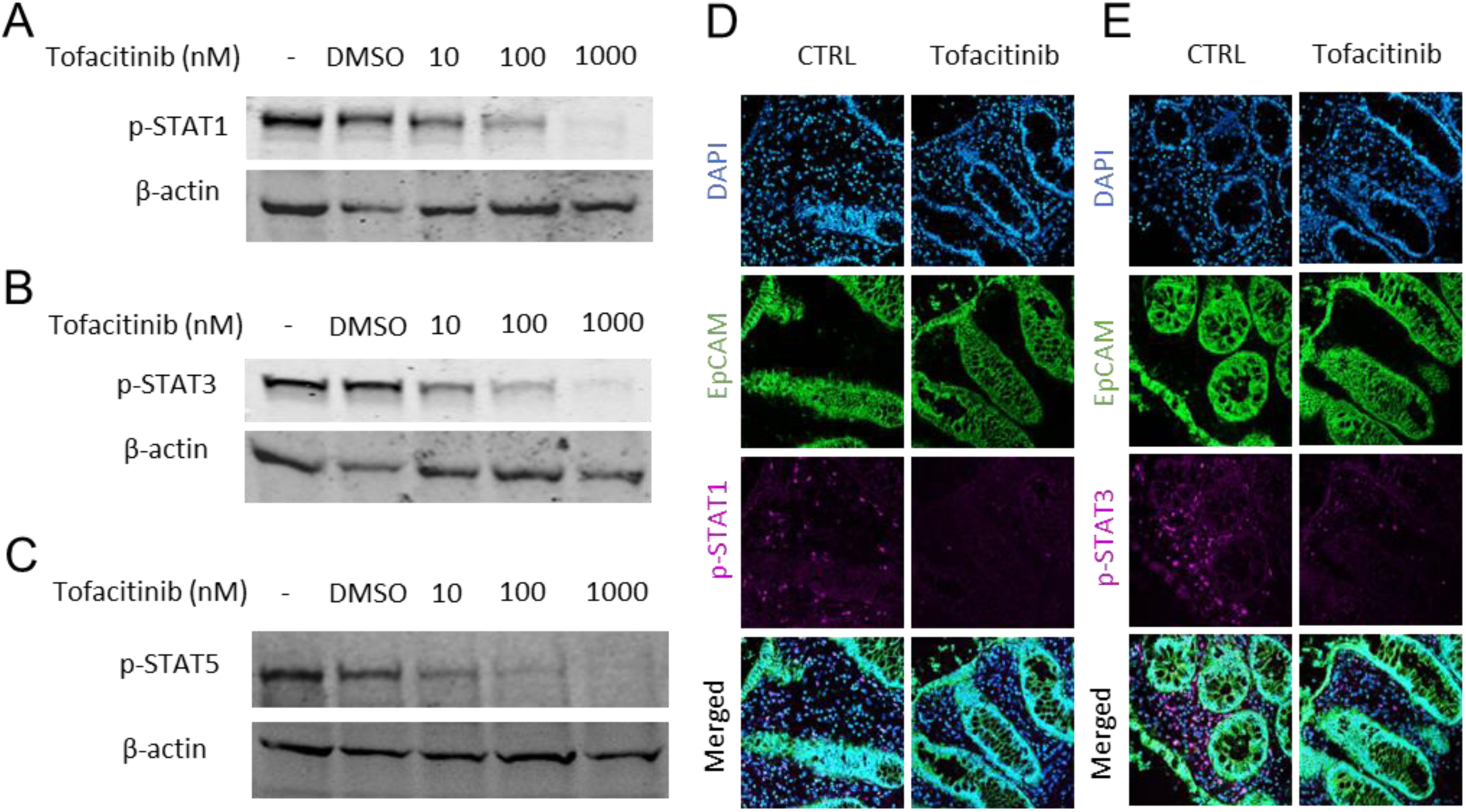
JAK inhibition decreases p-STATs expression in human mucosal explants. (A-C) Colonic mucosal explants from NC were treated with vehicle DMSO or with 10-1000nM of tofacitinib for 1.5h. p-STATs levels were assessed by Western blot of mucosal explants lysates using anti-p-STAT1 (Tyr701), anti-p-STAT3 (Tyr705), and anti-p-STAT5 (Tyr694), and compared to β-actin as a loading control (respectively). (D, E) Sections (10µm) of 100nM tofacitinib-treated colonic mucosal explants were stained with anti-p-STAT1 (Tyr701) and anti-p-STAT3 (Tyr705), anti-EpCAM and DAPI (respectively). Visualization by confocal microscopy. Original magnification x20. Shown are representative of 3 independent experiments with 3 colonic mucosal explants from NC. CTRL-non-treated control.

### Human intestinal organoids (HIOs) culture

HIOs were grown from crypts isolated from colonic NC surgical specimens. Crypt isolation and cell dissociation was based on Sato et al., 2011^13^. Briefly, Specimens were washed with sterile PBS, mucosa was stripped from the resected tissue, cut, and washed twice with crypt isolation medium (detailed in Supplementary Materials). Mucosa was incubated in a crypt isolation medium consisted of 0.5 mM DL-Dithiothreitol (DTT), 5.6 mM Na2HPO4, 8 mM KH2PO4, 96.2 mM NaCl, 1.6 mM KCl, 43.4 mM sucrose, 54.9 mM D-sorbitol supplemented with 2 Mm EDTA, for 30 minutes at 4°C followed by vigorously shaking. The crypt pellet was washed with FBS, resuspended in ice-cold Matrigel (FAL356231, Corning, New York, United States), and seeded as 15 µl domes on pre-warmed 12-well tissue culture plates. Plates were incubated upside down for 20 min in a 37°C 5% CO2 incubator until Matrigel solidification. Organoid expansion media was based on Pleguezuelos-Manzano et al., 2020^14^ and Usui et al., 2018^15^ and consisted of advanced DMEM: F12 (12634010, Gibco) (26% of total volume), 100 U/ml penicillin, 100 µg/ml streptomycin, 10 mM HEPES (10 mM, 03-025-1B, Biological Industries), 1× GlutaMAX (35050-038, Gibco), and the following growth factors: 1× B27(12587001, Gibco), 1 mM N-Acetylcysteine (A9165, Sigma-Aldrich), 100 ng/ml Noggin (120-10C, Peprotech), 50 ng/ml human EGF (AF-100-15, Peprotech), 10 mM Nicotinamide (N0636, Sigma-Aldrich), 10 μM SB202190 (1264, Tocris), 500 nM A83-01 (2939, Tocris), 10 nM Prostaglandine E2 (2296, Tocris), 26 μg/ml Primocin (ant-pm-1, InvivoGen), and conditioned medium from the L cell line secreting Wnt3A (50% of total volume) and 293T cells secreting R-spondin 1 (20% of total volume). 10 Μm Y-27632 (Y0503, Sigma-Aldrich, Missouri, United States) was added to the expansion medium for the first 2 days. The medium was changed every other day.

2D HIOs monolayer culture protocol was based on the supplementary protocol of Intesticult™ medium (WWW.STEMCELL.COM) culture. 96-well tissue culture plates were coated with 1:50 Matrigel in PBS for 1h at a 37°C 5% CO2 incubator. 3D HIOs were resuspended with Recombinant Trypsin EDTA Solution (03-079-1A, Biological Industries, Beit Haemek, Israel) and mechanically disrupted into a single-cell suspension. Cells were resuspended in an expansion medium containing 10Μm Y-27632 and seeded on Matrigel-coated plates (1×10^4^ cells/well). The expansion medium was changed every other day for 5-7 days. Prior to stimulations, HIOs were grown for additional 2-3 days in a generic differentiation medium based on Pleguezuelos-Manzano et al., 2020^14^, consisted of advanced DMEM F12 was supplemented with 100 U/ml penicillin, 100 µg/ml streptomycin, 10 mM HEPES, 1× GlutaMAX, 1× B27, 1 mM N-Acetylcysteine, 500 ng/ml human R-spondin 1 (120-38, Peprotech), 100 ng/ml Noggin, 50 ng/ml human EGF, and 100 μg/ml Primocin.

HIOs cultured for 7-12 days were treated with 0.1-10 µM indicated JAK inhibitor 1h prior to 50ng/ml human recombinant IFN-γ (300-02, PeproTech, Rehovot, Israel) stimuli for 24h. HIOs were harvested for Western blot and immunofluorescence as indicated below.

### RNA extraction

For RNA extraction, mucosal explants were homogenized in the ZR BashingBead Lysis Tube (S6012-50, Zymo Research, California, United States) using a high-speed bead beater (OMNI bead rupture 24). Total RNA was extracted from mucosal explants using Trizol® (15596026, Invitrogen, Massachusetts, United States) according to a standard protocol. RNA concentration and quality were assessed using NanoDrop Spectrophotometer (Thermo Scientific, Massachusetts, United States). All samples’ 260/280 and 260/230 ratios were >1.8.

### NanoString nCounter assay and statistical analysis

Gene expression in mucosal explants was analyzed using the nCounter Human Inflammation V2 Panel (NanoString, Washington, United States) according to the manufacturer’s instructions. Briefly, the hybridization buffer combined with the codeset of interest is combined with 5μl (400ng) of total RNA and incubated at 65°C overnight. Samples were then loaded onto the prep station and incubated under a high-sensitivity program for 3h. Following the prep station, samples were read using NanoString digital analyzer with the high-resolution option. To normalize the gene counts, the NanoString nSolver 4.0 software was used. The normalization was performed according to the standard protocol with thresholding according to the geometric mean of negative controls, normalization according to the geometric mean of positive controls, and the standard CodeSet housekeeping genes normalization (according to the geometric mean of GUSB, HPRT1, and TUBB genes). The normalized results were exported as a comma separated file (.csv) for further gene expression analysis in R version 4.1.0 (http://www.R-project.org/). Analysis of similarities test was done with *adonis* function in R package vegan. The differential expression between mucosal explants’ location (colon and ileum) or disease type (NC and UC, or NC and CD) was calculated using the Mann-Witney test. The differential expression between paired CTRL and tofacitinib-treated mucosal explants in each group (NC, UC, and CD) was calculated using Wilcoxon signed-rank test. Genes with a *P* value <0.05 were regarded as differentially expressed.

### Cell lysis and Western blot

Mucosal explants and HIOs were washed with ice-cold PBS and homogenized and solubilized, respectively, in lysis buffer (50 Mm HEPES Ph = 7.5, 150 Mm NaCl, 10% glycerol, 1% triton X, 1 Mm EDTA Ph = 8, 1 Mm EGTA Ph = 8, 1.5 Mm MgCl2), protease inhibitor cocktail set I (539131, Calbiochem, California, United States), and phosphatase inhibitor cocktail set II (524625, Millipore, Massachusetts, United States) and III (P0044, Sigma-Aldrich, Missouri, United States). Lysates were cleared by centrifugation. Protein concentration in the supernatant was measured using Bradford reagent (5000006, Bio-Rad, California, United States). Then, lysates were complemented with 4×sample buffer, heated at 95°C for 5 minutes, and stored at −20°C.

An equal amount (30µg) of cells lysates was subjected to electrophoresis on a sodium dodecyl sulfate (SDS) 7.5% or 4-20% polyacrylamide gel (4561024 or 4561095, respectively, Bio-Rad, California, United States). Proteins were semi-dry transferred to PVDF membranes (1704159SP5, Bio-Rad, California, United States) using Trans-Blot Turbo Transfer System (Bio-Rad, California, United States). Membranes were blocked for 1h in a Tris-Buffered Saline, 0.1% Tween® 20 (TBST, 2089232300, Bio-Lab, California, United States) buffer containing 6% nonfat milk, and reacted with the indicated primary antibodies, diluted in TBST supplemented with 5% BSA (82-045-1, Millipore, Massachusetts, United States), overnight at 4°C. Next, incubation with fluorescent conjugated secondary antibodies diluted in TBST was performed for 1h at room temperature. Signals were visualized using Odyssey CLx imager (Licor Biosciences, Nebraska, United States). Primary and secondary antibodies are detailed in Supplementary Table 1.

### Immunofluorescence

Frozen sections of mucosal explants and HIOs were fixed with 4% paraformaldehyde for 15 min at room temperature and washed with PBS. For permeabilization, frozen sections were incubated with absolute methanol for 10 min at −20°C and washed with PBS. Frozen sections and HIOs were next incubated in blocking buffer containing PBS, 5% normal donkey serum (017-000-121, Jacson ImmunoResearch, Pennsylvania, United States), and 0.3% TritonX100 (9036-19-5, Sigma-Aldrich, Missouri, United States) for 1h at room temperature and washed with PBS. Then, frozen sections and HIOs were stained with primary antibodies overnight at 4°C, followed by staining with fluorescence conjugated secondary antibodies for 1h at room temperature and counterstaining with DAPI, included in the mounting medium (GBI Labs). Frozen sections and HIOs were visualized by confocal microscope (LSM 800, Zeiss). Primary and secondary antibodies are detailed in Supplementary Table 1.

### Statistical analysis

Statistical analysis was performed with Prism version 9 (GraphPad Software). Data were checked for normality using Shapiro-Wilk test and then the appropriate test was applied. Differences were noted as significant by the following conventions: \**P*<0.05; \*\**P*<0.01; *** *P* <0.001 **** *P* <0.0001, as indicated. Bar graphs are shown as mean ± SD.

## Results

### Study population

The study included 68 patients, divided into two independent groups, depending on the source of mucosa and to enable specific analyses. In the first group, 26 mucosal explants were derived from colonic and ileal surgical resections from NC (n=13), patients with UC (n=3), and CD (n=13) and were used for the assessment of p-STATs expression levels. In the second group, 42 mucosal explants were derived from colonic and ileal biopsies from NC (n=15), patients with UC (n=17), and CD (n=10) and were used for the assessment of inflammatory gene expression levels. Patients’ clinical and demographic characteristics are detailed in Table 1.

### JAK inhibition decreases p-STATs expression in human mucosal explants

To elucidate JAK inhibition effect on signaling pathways in the human intestinal mucosa, tofacitinib-treated colonic mucosal explants from NC were analyzed for p-STATs. Colonic mucosal explants were treated with tofacitinib for 1.5h at different concentrations, and p-STATs were assessed by Western blot and immunofluorescence. A dose-dependent decrease in p-STAT1, p-STAT3, and p-STAT5 expression was observed in response to tofacitinib treated compared to non-treated control (CTRL) colonic mucosal explants from NC (Fig. 1A-C, respectively). Tofacitinib also potently inhibited p-STAT1 and p-STAT3, as assessed by immunofluorescence (Fig.1D and E, respectively). Notably, p-STATs expression was inhibited by tofacitinib but not by vehicle DMSO. The data indicate that intestinal mucosa could respond to *ex-vivo* JAK inhibition.

### Response to *ex-vivo* JAK inhibition is location and patient-dependent

To test whether JAK inhibition has a differential effect on STAT phosphorylation in colonic and ileal mucosa, or in the mucosa of patients with UC or CD, tofacitinib-treated mucosal explants were analyzed for p-STAT1, p-STAT3, and p-STAT5 (group 1; Table 1). Colonic or ileal mucosal explants obtained from NC, UC, and CD were treated with 100 nM tofacitinib for 1.5h, and p-STATs expression was assessed by Western blot. Tofacitinib potently inhibited p-STAT1 and p-STAT3 in colonic (Fig. 2A-C; 59.8 ± 17.1 and 50 ± 16.5 average inhibition rates ± SD, respectively) and ileal (Fig. 2C; 45.4 ± 13.8 and 41.9 ± 17.5 average inhibition rates ± SD, respectively) mucosal explants from NC. Furthermore, tofacitinib potently inhibited p-STAT1 and p-STAT3 in colonic mucosal explants from patients with UC (Fig. 2A and B; 50.1 ± 15.3 and 52.5 ± 18.4 average inhibition rates ± SD, respectively) and CD (Fig. 2A, B and D; 60.9 ± 18.2 and 52.6 ± 24.4 average inhibition rates ± SD, respectively), as wells as in ileal mucosal explants from patients with CD (Fig. 2D; 57 ± 20.2 and 38 ± 23.2 average inhibition rates ± SD, respectively). Variation in patients’ responses was observed, represented by the different rates of p-STATs inhibition, ranging between 10% to 88% (Fig. 2B-D). Disease-specific effects were not observed (Fig. 2B). Notably, p-STATs inhibition rates were higher in colonic compared to ileal mucosal explants (Fig. 2C, D). Similar results were observed when examining p-STAT5 (Fig. S1). Interestingly, p-STAT1 and p-STAT3 inhibition rates in colonic mucosal explants from patients with CD negatively correlated with C-reactive protein (CRP) levels (Fig. 2E; Pearson r= −0.82, *P*= 0.01 and Spearman r= −0.85, *P*= 0.02, respectively). No correlation was found between p-STATs inhibition rates and other patients’ characteristics (i.e., age or disease duration). In addition to reflecting the heterogeneity of patients with IBD, the results suggest that response to JAK inhibition, specifically by tofacitinib, is individual and location dependent. Thus, differential responsiveness between patients may be expected.

**Figure 2.**
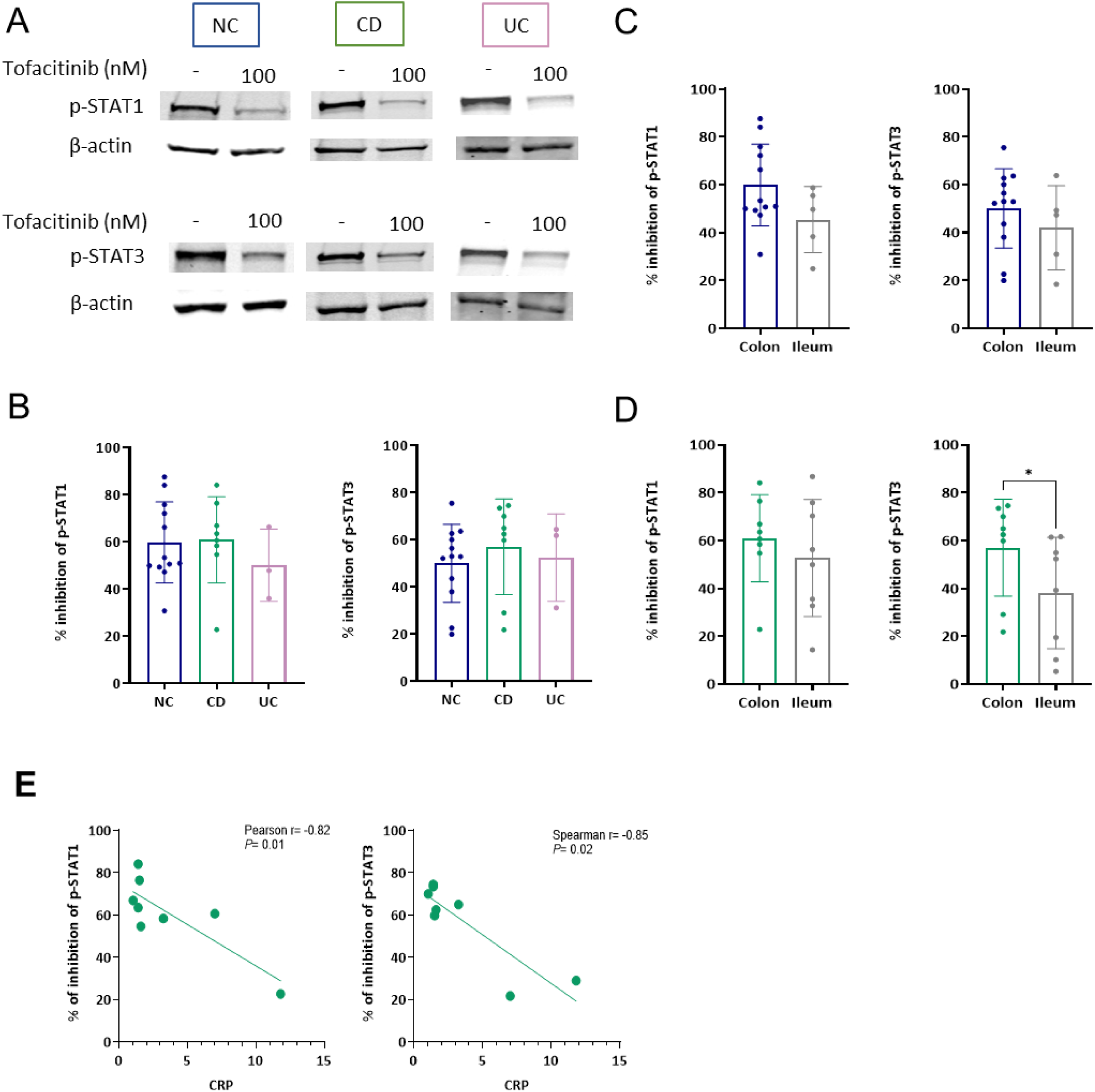
Response to *ex-vivo* JAK inhibition is location and patient dependent. Colonic and ileal mucosal explants from NC, patients with CD or UC were treated with 100nM of tofacitinib for 1.5h. (A) p-STATs levels were assessed by Western blot of colonic mucosal explants lysates using anti-p-STAT1and anti-p-STAT3 compared to β-actin as a loading control. (B-D) Densitometry analysis of the Western blot is represented as % of inhibition of p-STAT1 and p-STAT3 (ratio of p-STAT\β-actin) in 100nM tofacitinib-treated compared to the non-treated CTRL explant. (B) Colonic mucosal explants from NC, patients with UC or CD. (C, D) Colonic and ileal mucosa from NC and patients with CD, respectively. The results are presented as mean± SD. Statistical significance was calculated using paired t-test (* P < 0.05). (E) Correlation between % of inhibition of p-STAT1 (Pearson) and p-STAT3 (Spearman) in colonic tofacitinib-treated mucosal explants of patients with CD and CRP levels. NC-non-IBD control; UC-ulcerative colitis; CD-Crohn’s disease; CTRL-non-treated control; CRP-C-reactive protein.

### Intestinal mucosal explants have distinct inflammatory gene profiles according to location and disease phenotype

Next, we sought to assess the JAK inhibition effects downstream of STAT phosphorylation, thus examining alterations in gene transcription. To this end, we collected intestinal mucosal explants from NC and patients with UC and CD (group 2; Table 1). From each patient, mucosal explants were treated overnight with and without tofacitinib. Subsequently, mRNA expression levels of 255 inflammatory genes were assessed. To confirm that the mucosal explants maintain their inflammatory state after overnight culture, two unsupervised sample classifications were performed using principal component analysis (PCA). The first classification involved colonic and ileal mucosal explants from NC, while the second classification involved colonic mucosal explants from NC and patients with UC or CD. The PCA revealed a strong separation of colon and ileum from NC (analysis of similarities R2 = 0.31, *P* = 0.001), and of colonic mucosal explants from NC and patients with UC (analysis of similarities R2 = 0.32, *P* = 0.001) or CD (analysis of similarities R2 = 0.12, *P* = 0.044), based on their total inflammatory gene profiles (Fig. 3A, B and Fig. S2A, respectively). Additionally, colonic mucosal explants from NC demonstrated significant differential expressions of 179, 192, and 47 genes out of the 255 examined, when compared to the ileal mucosal explants from NC and to the colonic mucosal explants from patients with UC, and CD, respectively. Hierarchical clustering of these differentially expressed genes (DEGs) revealed a clear distinction between colonic and ileal mucosal explants from NC (Fig. 3C) and between colonic mucosal explants from NC and patients with UC or CD (Fig. 3D and Fig. S1B, respectively). The data indicates that mucosal explants cultured overnight exhibit unique inflammatory gene profiles depending on their original location and inflammatory state.

**Figure 3.**
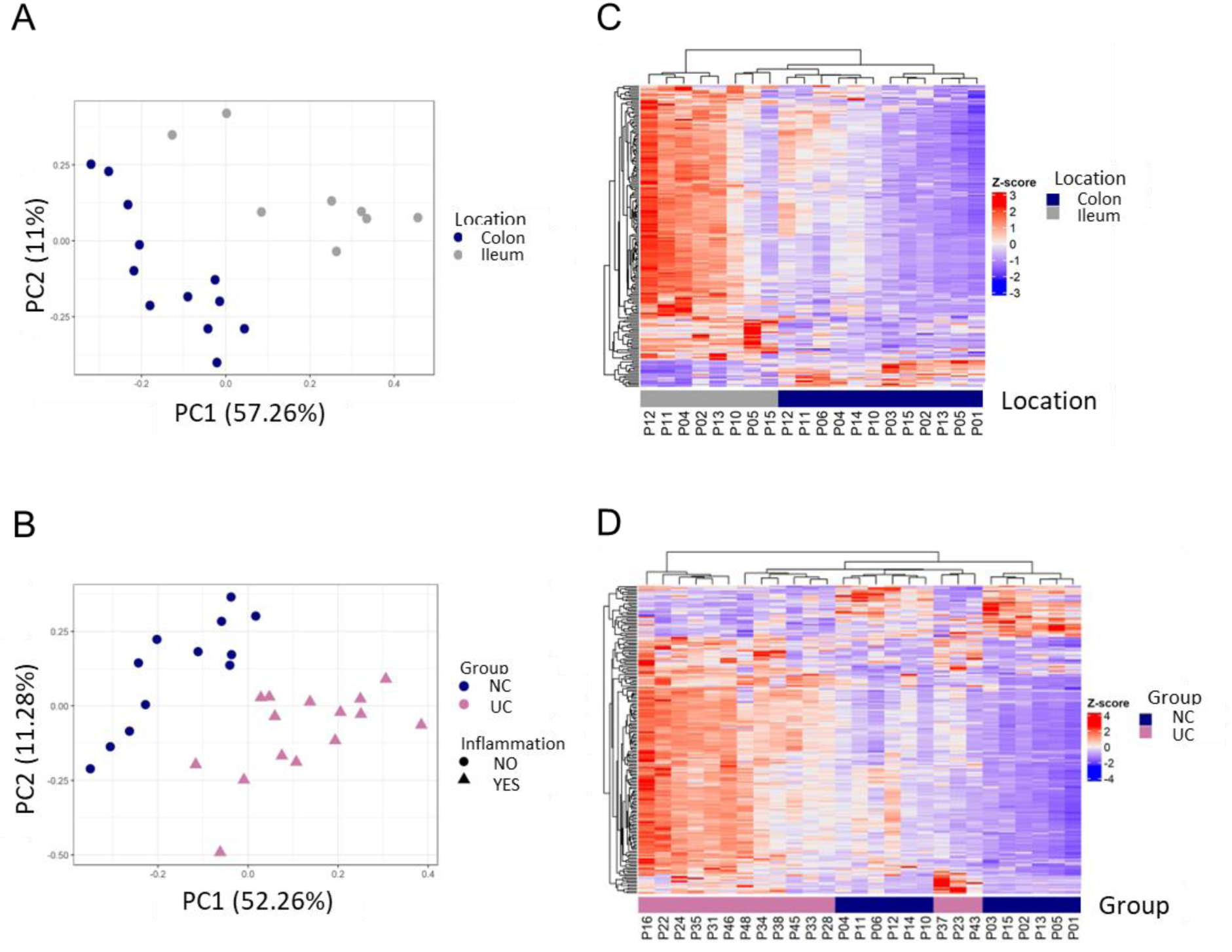
Intestinal mucosal explants have distinct inflammatory gene profiles according to location and disease phenotype. (A, B) PCA plots demonstrating 255 inflammatory genes expression profile of colonic and ileal mucosal explants from NC, and from colonic mucosal explants from NC and patients with UC cultured overnight, respectively. Each dot represents one sample, colored by group (NC in dark blue and UC in pink) and shaped by inflammation (noninflamed in circles and inflamed in triangles). Analysis of similarities separation test was performed for NC mucosal explants’ location (*R2* = 0.31, *P* = 0.001) and for colonic mucosal explants’ disease phenotype (*R2* = 0.32, *P* = 0.001). (C, D) Heatmap of the Z-score values of DEGs in ileal compared to colonic mucosal explants from NC, and in colonic mucosal explants from NC compared to colonic mucosal explants from patients with UC, respectively. Numbers below columns signify a patient number. NC-non-IBD control; UC-ulcerative colitis.

### A more robust response to JAK inhibition in colonic compared to ileal mucosal explants from NC

To identify molecular mechanisms underlying the anti-inflammatory response to JAK inhibition, we first examined JAK inhibition effects in the colon compared to the ileum in CTRL vs. tofacitinib-treated mucosal explants from NC (group 2; Table 1). Interestingly, PCA revealed a distinct separation between CTRL and tofacitinib-treated colonic mucosal explants, based on their total inflammatory gene expression profiles (analysis of similarities R2 = 0.12, *P* = 0.027, Fig. 4A). In contrast, no such separation was observed in ileal mucosal explants in response to the treatment (Fig. 4B). Furthermore, in response to JAK inhibition using tofacitinib, differential expression analysis in colonic mucosal explants revealed significant changes in 120 genes out of the 255 examined. Conversely, only 30 genes significantly changed after ileal mucosal explants were treated with tofacitinib. Hierarchal clustering of DEGs revealed relatively distinct gene expression alteration patterns in tofacitinib-treated compared to CTRL in both colonic and ileal mucosal explants (Fig. 4C and D, respectively). 17 DEGs were shared with colonic and ileal mucosal explants, including *STAT1, STAT2, STAT3, NOS2, and HIF1A* (Fig. 4E and F). These results further suggest a discordant, site-specific anti-inflammatory effect of JAK inhibition, specifically tofacitinib.

**Figure 4.**
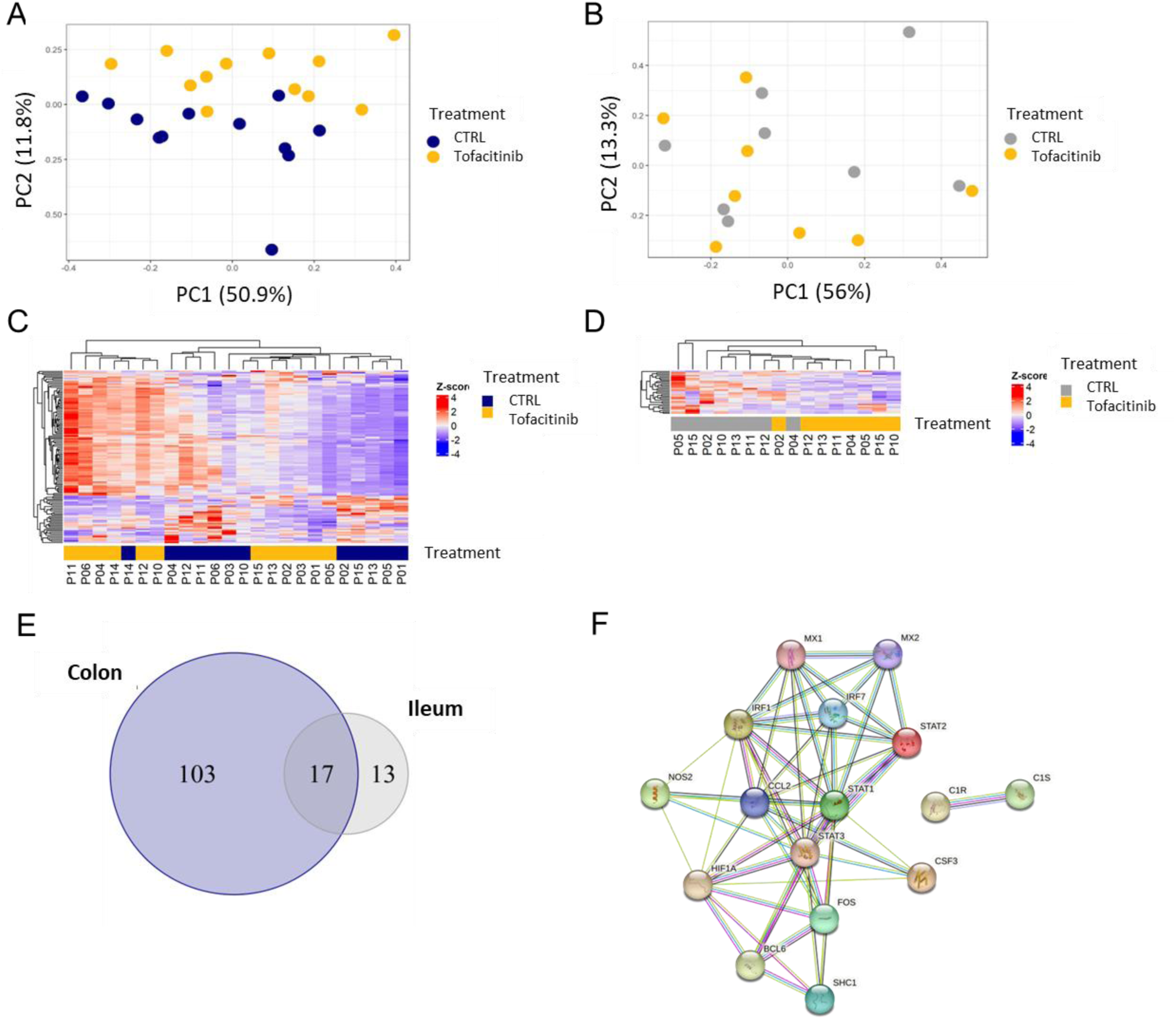
A more robust response to JAK inhibition in colonic compared to ileal mucosal explants from NC. Colonic and ileal mucosal explants from NC were CTRL or tofacitinib-treated overnight. (A, B) PCA plots of 255 inflammatory genes expression of colonic and ileal mucosal explants from NC, CTRL, or tofacitinib-treated overnight (respectively). Analysis of similarities separation test was performed for colonic and ileal mucosal explants according to treatment (*R2* = 0.12, *P* = 0.027 and *R*2= 0.06, *P* = 0.451, respectively). (C, D) Heatmaps of the Z-score values of DEGs in tofacitinib-treated compared to CTRL colonic and ileal mucosal explants, respectively. Numbers below the columns signify a patient number. (E) A Venn diagram showing the number of shared and unique DEGs in colonic and ileal mucosal explants in response to tofacitinib. (F) STRING network analysis. Networks visualizing the functional protein association for the 17 DEGs in both colonic and ileal mucosal explants from NC. Nodes in the network represent proteins. Edges represent protein–protein interactions, which (depending on the color) indicate known or predicted interactions. CTRL-vehicle DMSO control.

### JAK inhibition effects on inflammatory genes expression vary in colonic mucosal explants from patients with IBD

We next sought to assess JAK inhibition effects in CTRL compared to tofacitinib-treated mucosal explants from patients with IBD (group2; Table 1). PCA displayed no separation of CTRL and tofacitinib-treated colonic mucosal explants from patients with UC and CD based on their total inflammatory gene expression profiles (Fig. 5A and S3A, respectively). Additionally, while JAK inhibition in colonic mucosal explants from UC or CD significantly altered only five and six genes, respectively (Fig. 5B and S3B, respectively), none of the genes was significantly altered in ileal mucosal explants from patients with CD. Three DEGs were shared to colonic and ileal mucosal explants from NC and colonic mucosal explants from patients with UC; *STAT1, NOS2* and *MX1* (Fig. 5C). Interestingly, gene expression alteration patterns were variable among patients with IBD in response to JAK inhibition. For instance, while *STAT1*, *STAT2,* and *STAT3* expression levels decreased in all mucosal explants from NC, their expression increased in several colonic mucosal explants from patients with IBD (emphasized in red lines in Fig. 5D-F). This data further suggests that the *ex-vivo* response to JAK inhibition using tofacitinib is patient dependent and emphasizes the need for personalized medicine looking at patients as individuals.

**Figure 5.**
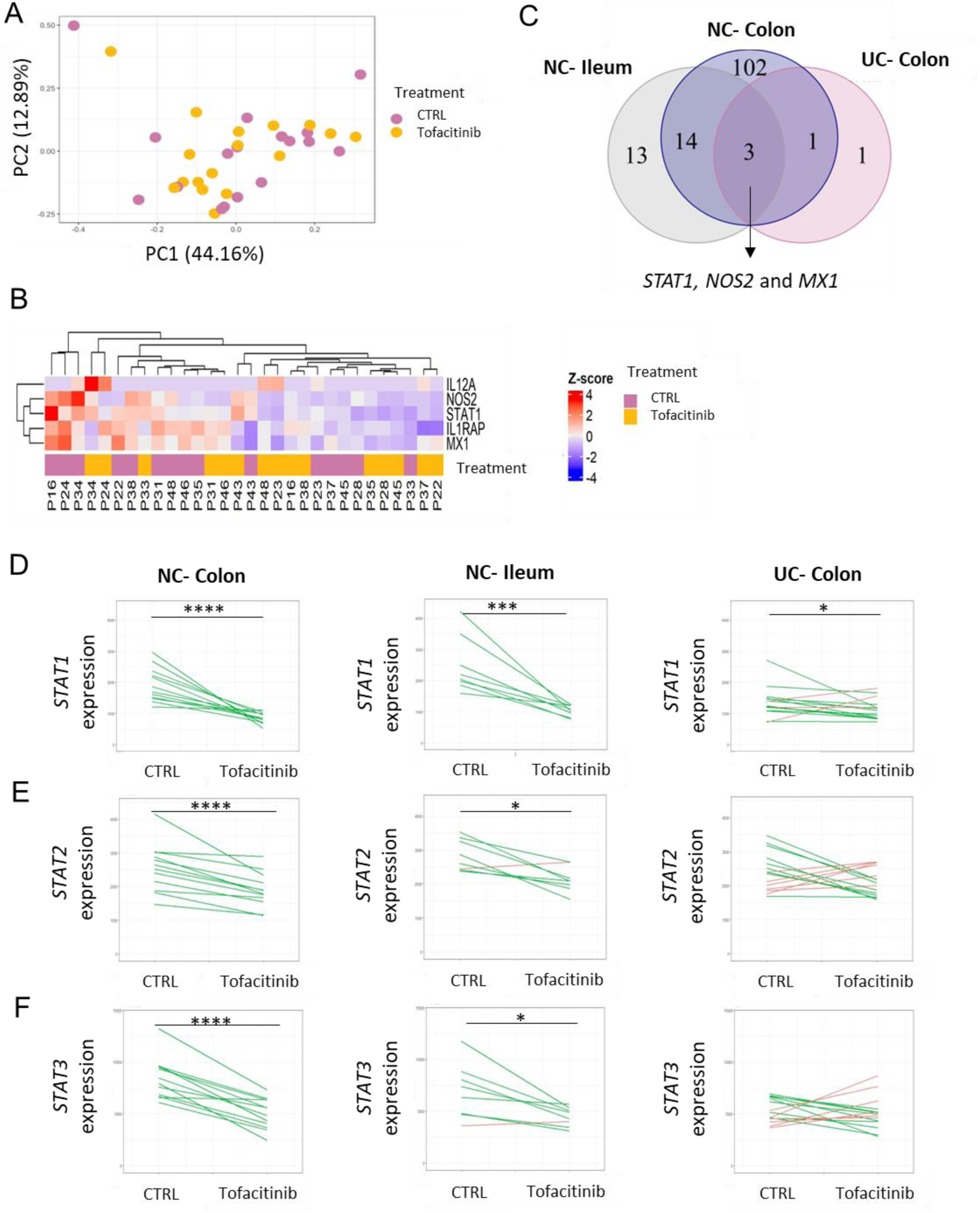
JAK inhibition effects on inflammatory genes expression vary in colonic mucosal explants from patients with IBD. (A) PCA plots of 255 inflammatory genes expression of colonic mucosal explants from patients with UC, CTRL or tofacitinib-treated overnight. (B) Heatmap of the scaled expression values of DEGs in tofacitinib-treated compared to CTRL colonic mucosal explants from patients with UC. Numbers below columns signify a patient number. (C) A Venn diagram showing the number of shared and unique DEGs in colonic and ileal mucosal explants from NC and colonic mucosal explants from patients with UC, in response to tofacitinib. (E, F and G) *STAT1, STAT2* and *STAT3* expression alterations in colonic and ileal mucosal explants from NC and colonic mucosal explants from patients with UC, CTRL or tofacitinib-treated overnight, respectively. Each line represents a patient. Reduced expression is in green and increased expression is in red (* *P* < 0.05, *** *P* < 0.001, **** *P* < 0.0001). NC-non-IBD control; UC-ulcerative colitis; CTRL-vehicle DMSO control.

### JAK inhibition decreases IFN-γ -induced p-STAT1 and inducible nitric oxide synthase (iNOS) upregulation in HIOs

*NOS2* was one of the shared DEGs to colonic and ileal mucosal explants from NC and colonic mucosal explants from patients with UC (Fig. 5C). *NOS2* encodes the iNOS, an enzyme responsible for producing nitric oxide that has been reported to be upregulated in the inflamed intestinal epithelium of patients with IBD^16–18^. To determine whether the decrease of *NOS2* in response to *ex-vivo* JAK inhibition is associated with decreased iNOS in response to JAK inhibition in an inflammatory condition, HIOs were treated with several JAK inhibitors of different specificities at increased concentrations 1 h before stimulation with IFN-γ for 24h. p-STAT1 and iNOS protein levels were assessed by Western blot and immunofluorescence. We examined the effect of the following JAK inhibitors: tofacitinib (reported inhibitory specificity JAK3 > JAK1\2), upadacitinib (JAK1), brepocitinib (JAK1\TYK2 >JAK2), and ritlecitinib (JAK3). Brepocitinib and ritlecitinib have shown efficacy in patients with UC in phase II clinical trials^19^ and their efficacy in patients with CD is currently evaluated^20^. Tofacitinib, upadacitinib, and brepocitinib significantly inhibited IFN-γ -induced p-STAT1 and iNOS expression levels in a dose-dependent manner (Fig. 6A, B, and C, respectively). No inhibition of p-STAT1 or iNOS expression levels was observed in response to ritlecitinib (Fig. 6D). IFN-γ -induced iNOS expression was completely blocked by the highest dose of JAK inhibitors; 10µM tofacitinib, upadacitinib, and brepocitinib, but not by ritlecitinib (Fig. 6A-E). These results further verify the downregulation of *NOS2* in response to JAK inhibition and underscore the potential of JAK inhibitors, such as tofacitinib, upadacitinib, and brepocitinib, to modulate iNOS expression in the context of intestinal inflammation.

**Figure 6.**
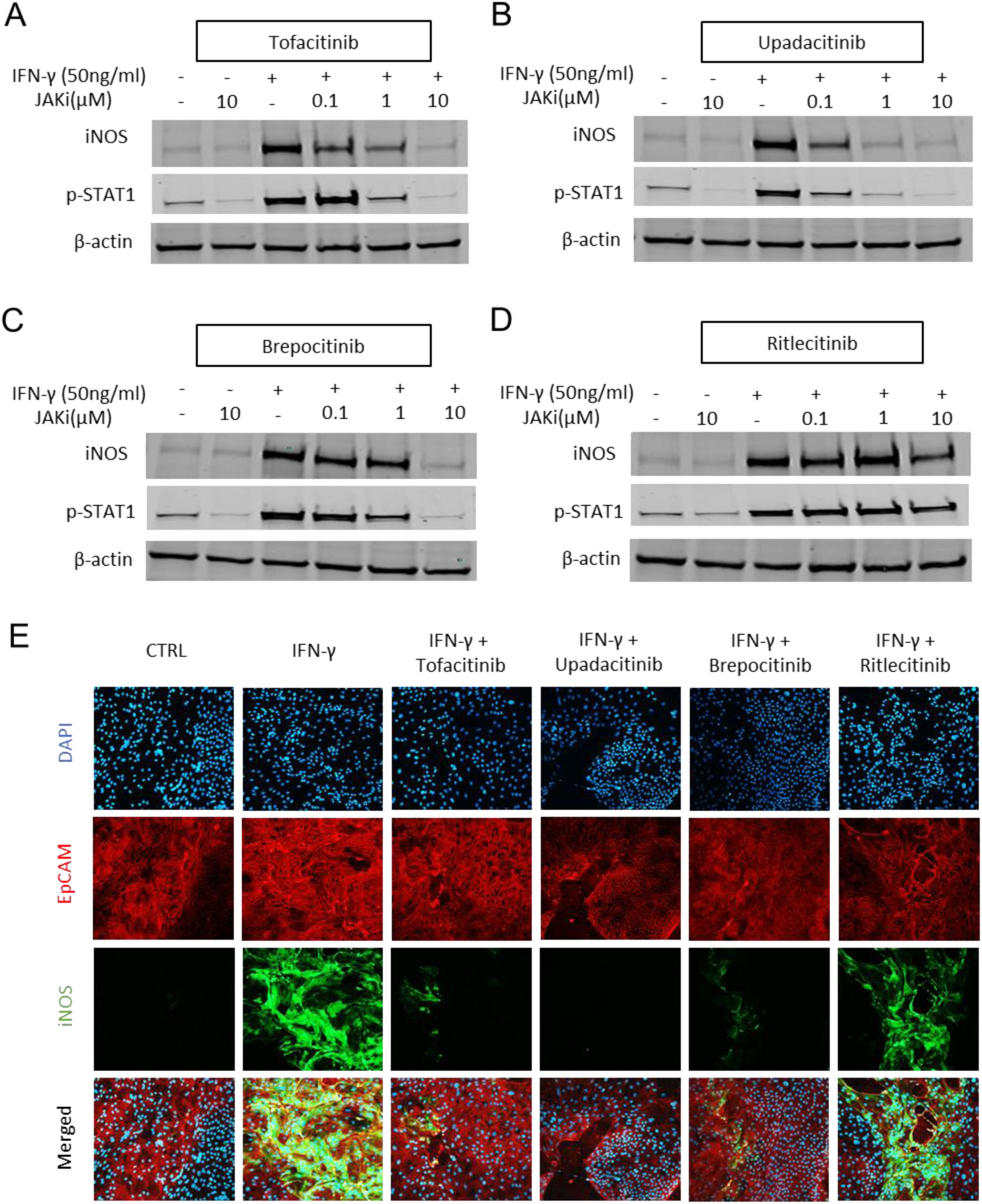
JAK inhibition decreases IFN-γ -induced p-STAT1 and iNOS upregulation in HIOs. (A, B, C, D) HIOs from NC were treated with vehicle DMSO control or increased concentrations (as stated) of tofacitinib, upadacitinib, brepocitinib, or ritlecitinib 1h before stimulation with 50ng/ml IFN-γ for 24h (respectively). p-STAT1 and iNOS levels were assessed by Western blot of HIOs lysates using anti-p-STAT1 and anti-iNOS, compared to β-actin as a loading control. (E) HIOs IFN-γ-treated with or without 10 µM of the indicated JAK inhibitors, were stained with anti-EpCAM, anti-iNOS, and DAPI. Visualization by confocal microscopy. Original magnification x10.Shown are representative of three independent experiments (n=3). CTRL-non-treated control; JAKi-JAK inhibitor.

## Discussion

JAK inhibitors used for the treatment of IBD, inhibit JAK-STAT signaling which transduces multiple pro-inflammatory cytokine signals. As with other IBD treatments, some patients do not respond to JAK inhibitors, and response to specific JAK inhibitors may vary. For instance, in real-world data of tofacitinib’s effectiveness in the treatment of patients with UC, clinical remission rates in week 8 ranged between 31-57%^21–23^. Among patients followed until week 52, clinical remission was achieved by only 41% of patients with UC receiving tofacitinib^23,24^ and by 52% of patients with UC receiving upadacitinib^11^, indicating individual patients’ responsiveness. Investigation of JAK inhibition effects on the entire human intestinal mucosa has been scarce^25,26^and the process and responses to JAK inhibitors in the intestinal mucosa remain elusive. In this study, we compared the *ex-vivo* mucosal response to JAK inhibition of patients with different IBD phenotypes and locations. Our experimental approach was designed to analyze the molecular changes occurring in the human intestinal mucosa in response to JAK inhibition and to assess individual responses.

We demonstrated that overnight-cultured colonic and ileal mucosal explants from NC or patients with IBD have distinct constitutive inflammatory gene profiles according to location and disease phenotype. While prior research showed higher secretion levels of cytokines in overnight-cultured samples from UC compared to samples from NC^25^, our study further elucidated that the differences extend beyond cytokine levels. We observed significant differences in multiple inflammation-related genes in overnight-cultured mucosal explants from patients with IBD compared to NC. Consequently, our findings affirm the capacity of the human mucosal explants platform to accurately reflect an inflammatory state of the intestinal mucosa.

Using human mucosal explants, we showed that JAK inhibition using tofacitinib significantly alters more inflammatory genes and STAT phosphorylation in the colonic compared to ileal mucosal explants. Our results suggest that JAK inhibition, specifically by tofacitinib, may exert a stronger anti-inflammatory effect in the colon than in the ileum. While tofacitinib has been proven to be efficacious in patients with UC^21–24^, clinical trials examining the effect of tofacitinib in patients with CD failed to achieve the primary endpoints^9,10^. Notably, the clinical trials in CD included patients with ileal, colonic, and ileocolonic disease, without subgroup analysis examining the effects of tofacitinib according to disease location. Our results offer an explanation for the discordant response observed in the reported clinical efficacy of tofacitinib in patients with UC and CD. In recent years there is increasing evidence that ileal CD differs from colonic CD in several aspects, including the response to therapies. Higher response rates in colonic compared to ileal disease were reported in patients with CD receiving anti-TNFα agents, vedolizumab, or ustekinumab^27–30^. In contrast, lower response rates to enteral nutrition in colonic CD compared to ileal CD were reported^31,32^. While JAK1 inhibitors, including filgotinib and upadacitinib, have been shown to induce clinical remission in patients with CD^12,33^, a recent proof-of -concept study focusing on patient with small bowel Crohn’s disease, filgotinib did not show statistically significant differences vs placebo in the proportion of patients who achieved clinical remission^34^.Moreover, in a meta-analysis examining the molecular differences between colonic and ileal IBD at the transcriptomic level, colonic samples of patients with CD resembled those of patients with UC and differed from ileal samples of patients with CD^35^. Overall, the molecular mechanisms underlying IBD in the colon may be different from those in the ileum, affecting response to therapy. Our study highlights that this distinction may also extend to the effects of JAK inhibitors. Nevertheless, whether JAK inhibition has a differential effect in colonic and ileal disease *in-vivo* is unknown. Therefore, further analysis of JAK inhibition in patients stratified according to colonic or ileal location is needed in order to confirm our findings, enable personalized approach, and determine the mechanisms underlying these effects.

Another important finding is the variable, individual *ex-vivo* response to JAK inhibition using tofacitinib. We observed distinct patterns of altered gene expression, specifically in *STAT1*, *STAT2*, and *STAT3*, in mucosal explants from patients with IBD exposed to the JAK inhibitor tofacitinib. This suggests that individual patients may exhibit unique responses to JAK inhibition, potentially influencing treatment efficacy. Our results also demonstrate that the inhibition rates of p-STAT1, p-STAT3, and p-STAT5 varied widely, ranging from 10% to 88%. This variability in response to *ex-vivo* JAK inhibition further emphasizes the complex and heterogeneous nature of IBD, which is influenced not only by disease location but also by factors such as patients’ genetics, microbiome composition, nutrition, and immune system functionality^36^. It is evident that personalized and tailored approaches to IBD treatment are crucial to address this inherent heterogeneity^37–39^. By studying mucosal explants, we were able to capture and assess individual effects of JAK inhibition, specifically tofacitinib, on gene expression and pathway alterations. Therefore, our findings underscore the relevance of studying *ex-vivo* responses to predict patients’ responses, further supporting the exploration of personalized medicine approaches in the treatment of IBD.

Of note, colonic mucosal explants from CD also responded to tofacitinib–related JAK inhibition. JAK inhibition by tofacitinib led to reduced p-STAT1, p-STAT3, and p-STAT5 expression (Fig 2A, B and Fig S1D), along with downregulation of inflammatory pathway genes (Fig S3). These findings imply individual beneficial effects of tofacitinib in this population as well. Thus, re-evaluating pivotal tofacitinib studies focusing on patients with colonic disease is suggested, as well as prospective stratification of patients treated with JAK inhibitors according to disease location.

An interesting negative correlation emerged between the inhibition rates of p-STAT1 and p-STAT3 in tofacitinib-treated compared to CTRL mucosal explants from patients with CD and their levels of CRP, a well-established biomarker of inflammation, indicating that patients with higher baseline CRP levels exhibited reduced responsiveness to *ex-vivo* tofacitinib treatment. This finding aligns with previous *post-hoc* analyses of clinical trials such as OCTAVE Sustain, indicating that high CRP at baseline was associated with a reduced likelihood of remission at week 52^40^. Moreover, a real-world study reported similar results, indicating that higher CRP levels at baseline were associated with primary non-response to tofacitinib (*P*=0.004)^21^. Therefore, monitoring CRP levels could potentially serve as a useful biomarker contributing to the prediction of patient responses to JAK inhibitors.

We identified *NOS2* as a significantly inhibited gene by tofacitinib. This significant inhibition was shared in colonic and ileal mucosal explants from NC and colonic mucosal explants from UC. To demonstrate the engagement of downstream targeted molecule of JAK inhibition, iNOS was chosen as a candidate molecule to be further assessed in HIOs. iNOS and the downstream nitric oxide expression are increased in the mucosa of patients with IBD compared to NC^41,42^, located in the intestinal crypts and epithelial cells^16,18^. Excessive amounts of nitric oxide may act as a pro-inflammatory mediator, leading to oxidative stress and tissue damage and exacerbating the inflammatory response in the intestinal mucosa^43^. While a previous study highlighted the importance of JAK-STAT signaling in IFN/LPS-induced iNOS in rat intestinal epithelial cell line^44^, we demonstrated that JAK inhibition, using several JAK inhibitors, decreases iNOS expression in human intestinal epithelial cells. Moreover, our findings suggest that the extent of IFN-γ -induced iNOS reduction depends on the specificity of the JAK inhibitor used. These findings provide valuable evidence for the anti-inflammatory effects associated with JAK inhibitors within the intestinal mucosa, suggesting their potential utility in ameliorating oxidative stress and tissue damage in patients with IBD.

Our study has several notable strengths. The study was conducted within a single, well-established tertiary IBD center, allowing us to recruit a substantial number of patients. Furthermore, we were able to correlate our *ex-vivo* findings with patients’ characteristics and clinical data. Experiments were done using freshly obtained human intestinal mucosa, thus having immediate, practical relevance to patients. Remarkably, we observed an intriguing correlation between the *ex-vivo* JAK inhibition effect and patients’ CRP expression levels, mirroring the reported correlation between clinical response to JAK inhibition and CRP levels. This provided further supporting evidence for the clinical relevance of our *ex-vivo* platform. Finally, the translational, multidisciplinary approach enabled deciphering molecular pathways relevant to JAK inhibitors effects, adding knowledge regarding their mechanism of action in the human intestinal mucosa. Overall, using our *ex-vivo* platform we provide unique mechanistic insights, as well as meaningful clinical and translational implications of highest relevance for patient care.

While our study offers valuable insights, we acknowledge several limitations. The *ex-vivo* platform itself has several limitations, such as tissue dissociation from the blood flow, nervous system, and immune cells trafficking. Nonetheless, this model system mimics the tissue complexity and is the closest we can get to testing drugs on human intestinal mucosa without medical intervention. Additionally, a small number of ileal biopsies were obtained from patients with CD, limiting the analysis of inflammatory gene expression within this subgroup.

In conclusion, by studying multiple disease types and locations, we found that JAK inhibitors may have distinct anti-inflammatory effects in specific areas of the intestinal mucosa, with a stronger effect observed in the colon compared to the ileum. Furthermore, we found considerable interpatient variability in the *ex-vivo* response to JAK inhibition, suggesting individual patients’ responses and emphasizing that personalized treatment approaches are crucial in IBD management. Additionally, JAK inhibitors may have further therapeutic benefits by downregulating iNOS expression. *Ex-vivo* mucosal platforms may be a valuable resource for studying drugs effects and for personalized treatment strategies based on individual patient mucosal responses.

## Abbreviations

CD: Crohn’s disease
CRP: C-reactive protein
CTRL: Control
DEGs: Differentially expressed genes
HIOs: Human intestinal organoids
IBD: Inflammatory bowel diseases
IFN: Interferon
IL: Interleukin
iNOS: Inducible nitric oxide synthase
JAK: Janus kinase
JAKi: Janus kinase inhibitor
NC: Non-IBD controls
NOS2: Nitric oxide synthase 2
PCA: Principal component analysis
p-STAT: Phosphorylated signal transducers and activators of transcription
STAT: Signal transducers and activators of transcription
TNF: Tumor necrosis factor
TYK: Tyrosine kinases
UC: Ulcerative colitis

## Data Transparency Statement

Raw data and analytic methods will be made available to other researchers from the corresponding author upon request.

## Fundings

This work was partially supported by a generous grant from The Leona M. and Harry B. Helmsley Charitable Trust to Iris Dotan and an investigator-initiated research grant from Pfizer to Keren M Rabinowitz. The funders had no role in study design, data collection and analysis, decision to publish, or manuscript preparation.

## Notes

### Competing Interest Statement

KK, HAT, JA, ESB, NW, LHM, AF, ALB, ZL, HBE, IAB, SCK and KMR declare no conflict of interests. IG reports: research support from Gilead, Boehringer Ingelheim, Pfizer, and AbbVie. HY reports: institutional research grants from Pfizer and the ISF; consulting fees from AbbVie, Janssen, Pfizer, Takeda, Bristol Myers Squibb, and Elly Lilli; honoraria for lectures from AbbVie, Janssen, Pfizer, and Takeda; participation in a Data Safety Monitoring Board or Advisory Board for AbbVie, Pfizer, Takeda, Bristol Myers Squibb, and Elly Lilli. JEO reports: speaker fees from Takeda, Pfizer, Jansen, AbbVie, and Novartis; grant support from Pfizer; participation in Advisory Board of Takeda. ID has served as a speaker, consultant, and advisory board member AbbVie, Altman Research, Athos, Celltrion, Celgene/BMS, Ferring, Eli-Lilly, Gilead, Galapagos, Gutreat, Harp Diagnostics, Iterative Scopes, Janssen, Pfizer, Roche/Genentech, Sangamo, Sublimity, Sandoz, Takeda, Prometheus

