## Supplementary Table 1 for "Discordant effects of *ex-vivo* JAK inhibition on inflammatory responses in colonic compared to ileal mucosa"

| Antibody | Host | Dilution WB | Dilution IF | Cat. number | Company |
|---|---|---|---|---|---|
| **p-STAT1 (Tyr701)** | Rabbit monoclonal | 1:1000 | 1:200 | 9167 | Cell signaling |
| **p-STAT3 (Tyr705)** | Rabbit monoclonal | 1:2000 | 1:200 | 9145 | Cell signaling |
| **p-STAT5 (Tyr694)** | Rabbit monoclonal | 1:1000 | - | 4322 | Cell signaling |
| **p-STAT6 (Y641)** | Rabbit monoclonal | 1:1000 | - | 56554 | Cell signaling |
| **iNOS** | Rabbit monoclonal | 1:1000 | 1:800 | 20609 | Cell signaling |
| **α-tubulin** | Mouse monoclonal | 1:5000 | - | Ab7291 | Abcam |
| **β-actin** | Rabbit polyclonal | 1:2000 | - | Ab8227 | Abcam |
| **Anti-mouse IRDye680** | Goat | 1:10000 | | Ab216776 | Abcam |
| **Anti-rabbit IRDye800** | Goat | 1:10000 | | Ab216773 | Abcam |
| **Anti-mouse AF488** | Donkey | - | 1:1000 | Ab150109 | Abcam |
| **Anti-mouse AF647** | Donkey | - | 1:1000 | Ab150107 | Abcam |

Supplementary Table 1. Antibodies used for Western blot and immunofluorescence. WB- Western blot; IF- Immunofluorescence; p-STAT- Phosphorylated signal transducers and activators of transcription; iNOS- inducible nitric oxide synthase
