## supplementary figure 1 for "Discordant effects of *ex-vivo* JAK inhibition on inflammatory responses in colonic compared to ileal mucosa"

Figure S1

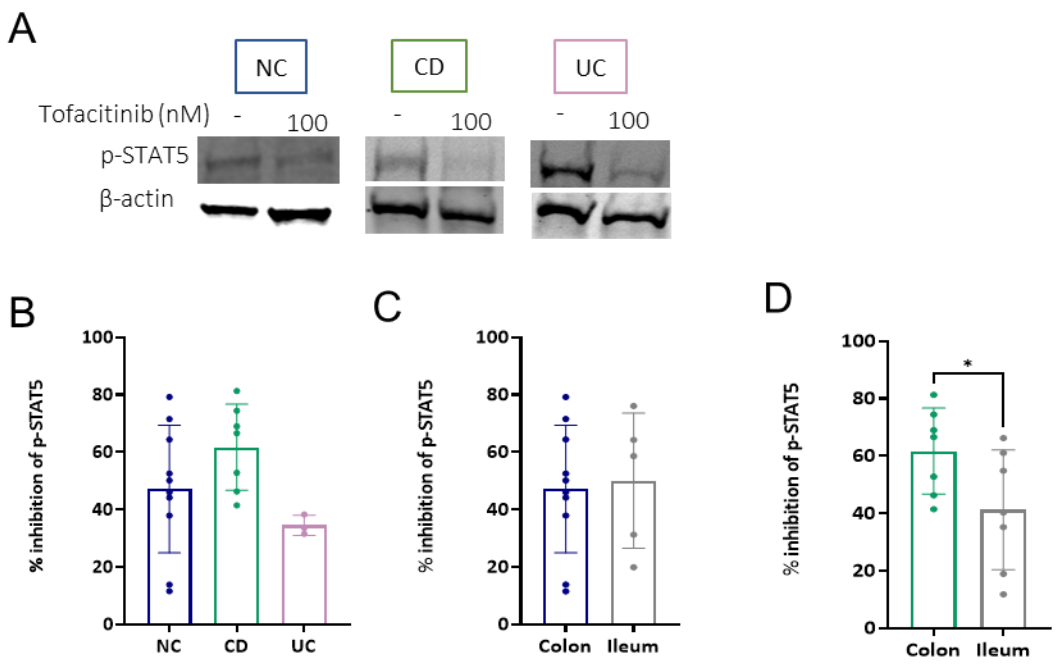


Figure S1. *Ex-vivo* JAK inhibition of p-STAT5 is location and patient-dependent. Colonic and ileal explants from NC, patients with CD or UC were treated with 100nM of tofacitinib for 1.5h. (A) p-STAT5 levels were assessed by Western blot of colonic explants lysates using anti-p-STAT5 compared to β-actin as a loading control. (B-D) Densitometry analysis of the Western blot is represented as % of inhibition of p-STAT5 (ratio of p-STAT5\β-actin) in 100nM tofacitinib-treated compared to the non-treated CTRL explant. (B) Colonic explants from NC, patients with UC or CD. (C, D) Colonic and ileal mucosa from NC and patients with CD, respectively. The results are presented as mean± SD. Statistical significance was calculated using paired t-test (* P < 0.05). NC-non-IBD control; UC- ulcerative colitis; CD-Crohn's disease; CTRL-non-treated control.
