## supplementary figure 2 for "Discordant effects of *ex-vivo* JAK inhibition on inflammatory responses in colonic compared to ileal mucosa"

Figure S2

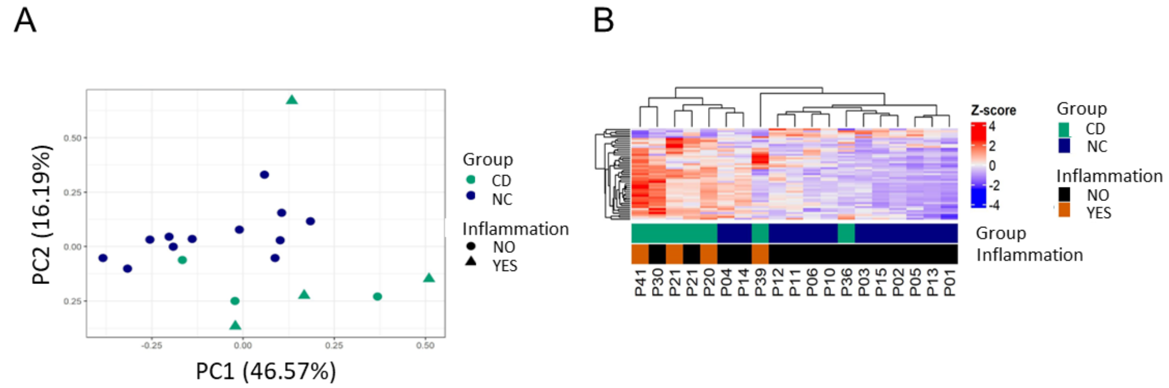


Figure S2. Intestinal explants have distinct inflammatory gene profiles according to disease phenotype. (A) PCA plots demonstrating 255 inflammatory genes expression profile of colonic explants from NC and patients with CD cultured overnight, respectively. Each dot represents one sample, colored by group (NC in dark blue and CD in green) and shaped by inflammation (noninflamed in circles and inflamed in triangles). Analysis of similarities separation test was performed for colonic explants' disease phenotype (*R2* = 0.12, *P* = 0.044). (B) Heatmap of the Z-score values of DEGs in colonic explants from NC compared to colonic explants from patients with CD. Numbers below columns signify a patient number. NC-non-IBD control; CD-Crohn's disease; DEGs- Differentially expressed gene.
