## supplementary figure 3 for "Discordant effects of *ex-vivo* JAK inhibition on inflammatory responses in colonic compared to ileal mucosa"

Figure S3

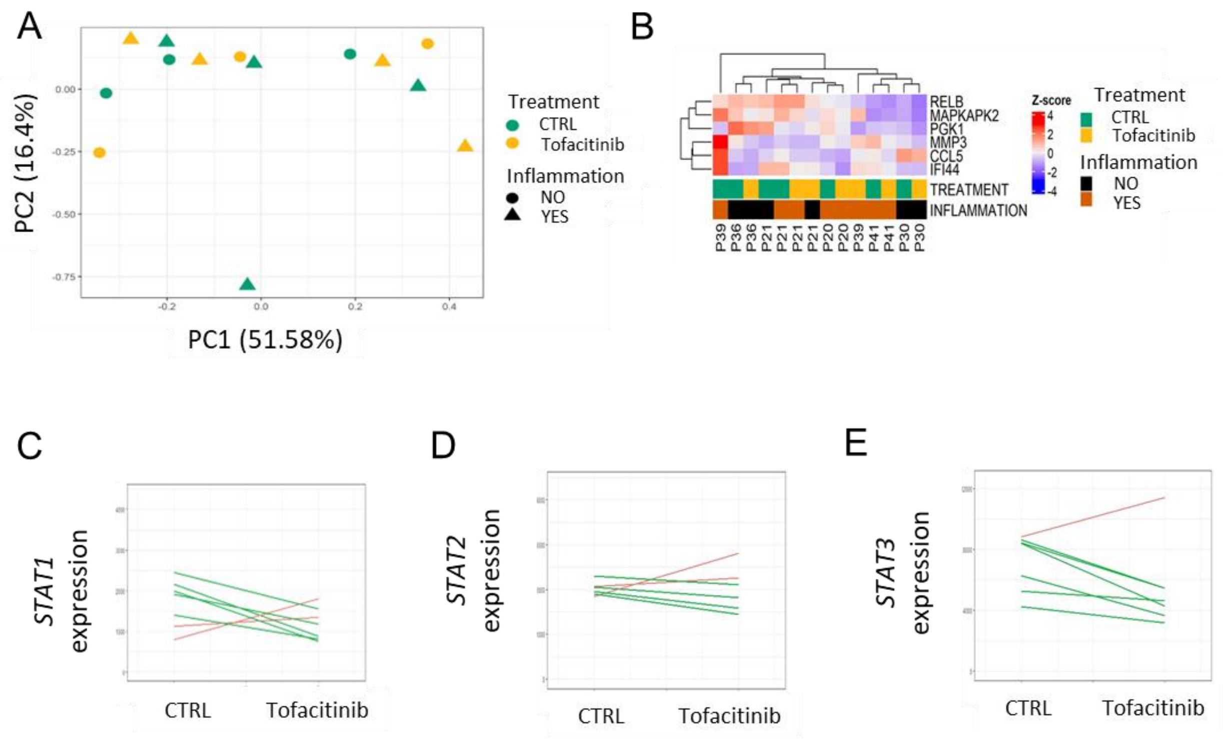


Figure S3. JAK inhibition effects on Inflammatory genes expression vary in colonic explants from patients with IBD. (A) PCA plots of 255 inflammatory genes expression of colonic explants from patients with CD, CTRL or tofacitinib- treated overnight. (B) Heatmap of the scaled expression values of DEGs in tofacitinib- treated compared to CTRL colonic explants from patients with CD. Numbers below columns signify a patient number. (C, D, and E) *STAT1, STAT2,* and *STAT3* expression alterations in colonic explants from patients with CD, CTRL or tofacitinib- treated overnight, respectively. Each line represents a patient. Reduced expression is in green and increased expression is in red. CD- Crohn’s disease; CTRL- vehicle DMSO control.
